# Staufen 1 amplifies pro-apoptotic activation of the unfolded protein response

**DOI:** 10.1101/820225

**Authors:** Mandi Gandelman, Warunee Dansithong, Karla P Figueroa, Sharan Paul, Daniel R Scoles, Stefan M Pulst

**Affiliations:** Department of Neurology, University of Utah, 175 North Medical Drive East, 5th Floor, Salt Lake City, Utah 84132, USA

## Abstract

Staufen-1 (STAU1) is an RNA binding protein that becomes highly overabundant in numerous neurodegenerative disease models, including those carrying mutations in presenilin1 (PSEN1), microtubule associated protein tau (*MAPT*), huntingtin (*HTT*), TAR DNA-binding protein-43 gene (*TARDBP*) or C9orf72. We previously reported that elevations in STAU1 determine autophagy defects. Additional functional consequences of STAU1 overabundance, however, have not been investigated. We studied the role of STAU1 in the chronic activation of the Unfolded Protein Response (UPR), a common feature among the neurodegenerative diseases where STAU1 is increased, and is directly associated with neuronal death. Here we report that STAU1 is a novel modulator of the UPR, and is required for apoptosis induced by activation of the PERK-CHOP pathway. STAU1 levels increased in response to multiple ER stressors and exogenous expression of STAU1 was sufficient to cause apoptosis through the PERK-CHOP pathway of the UPR. Cortical neurons and skin fibroblasts derived from *Stau1^−/−^* mice showed reduced UPR and apoptosis when challenged with thapsigargin. In fibroblasts from SCA2 patients or with ALS-causing TDP-43 and C9ORF72 mutations we found highly increased STAU1 and CHOP levels in basal conditions. STAU1 knockdown restored CHOP levels to normal. Taken together, these results show STAU1 overabundance reduces cellular resistance to ER stress and precipitates apoptosis.

## Introduction

Staufen-1 (STAU1) is a RNA-binding protein that localizes to stress granules during stress and can shape a cell’s transcriptome through multiple mechanisms, including regulation of translation efficiency, stress granule assembly, mRNA transport and Staufen-mediated mRNA decay (SMD) (1–11). We recently identified STAU1 as an interactor of wildtype and mutant ATXN2, which causes the polyglutamine disease spinocerebellar ataxia type 2 (SCA2) (2). We subsequently discovered substantial increases in STAU1 in multiple cell and animal models of human neurodegenerative diseases, including those carrying mutations in presenilin1 (PSEN1), microtubule associated protein tau (*MAPT*), huntingtin (HTT), TAR DNA-binding protein-43 gene (*TARDBP*) or C9orf72-SMCR8 complex subunit (*C9orf72*) (1), as well as stroke and myotonic dystrophy (3, 12). STAU1 overabundance can also be triggered by a variety of acute noxa such as calcium increase, ER stress, hyperthermia and oxidative stress (1, 13) Autophagy defects and IRES-mediated translation are the currently described mechanisms for STAU1 increase, both relevant protein abundance regulatory pathways during neurodegeneration (1, 13).

In neurodegenerative diseases, pathological mutations cause an increased load of misfolded and aggregated proteins and alterations in calcium homeostasis, leading to ER stress and activating the unfolded protein response (UPR) (14, 15). The UPR is a coordinated cellular response orchestrated by 3 main signaling pathways downstream of protein kinase RNA-like ER kinase (PERK), inositol-requiring enzyme 1 (IRE1) and activation transcription factor 6 (ATF6) (15). Activation of the UPR upregulates adaptive mechanisms that promote proper protein folding and regulate calcium balance, including chaperone gene expression, global suppression of protein synthesis and stimulation of autophagy and the proteasome. Failure to restore ER homeostasis leads to eIF2α phosphorylation downstream of PERK and induction of the pro-apoptotic transcription factor C/EBP homologous protein (CHOP), triggering the intrinsic apoptotic pathway (15–17). In addition of signaling through the UPR, phosphorylation of eIF2α is a critical early step of stress granule formation, as inhibition of protein synthesis leads to the aggregation of inactive translation complexes into stress granules (18, 19).

Here we demonstrate that STAU1 is required for the activation of apoptosis triggered by ER stress. Accordingly, STAU1 knockout cells were refractory to apoptosis induced by ER stress. STAU1 knockdown was sufficient to prevent the terminal activation of the UPR in cellular and animal models of SCA2 and ALS associated with improvement of motor deficits *in vivo* (2). In all, our study describes a novel connection between the RNA-granule protein STAU1 and ER stress-induced apoptosis that can be targeted in neurological diseases.

## Methods

### Cell lines and cell culture

Cell culture media and reagents were purchased from ThermoFisher Scientific unless specified. HEK293 cells and fibroblasts were maintained in DMEM supplemented with 10% fetal bovine serum (FBS). Gene editing of endogenous ATXN2 in HEK293 cells to express ATXN2 with 58 CAG repeats was performed with CRISPR/Cas9 according to published protocols (20), as detailed previously in our work (2). Cells were periodically screened by PCR to confirm the preservation of ATXN2-Q58. Fibroblasts from patients were obtained from a skin punch biopsy or from Coriell Cell Repositories (Camden, NJ, USA). All subjects biopsied gave written consent and procedures were approved by the Institutional Review Board at the University of Utah. Supplementary table 1 lists all human fibroblasts, their genetic mutation and repository identification number. All mutations were verified by PCR sequencing. Identity authentication of HEK293 cells and human fibroblasts was carried out by short tandem repeat (STR) analysis with the GenePrint 24 System (Promega, USA).

### Mice

All mice were housed and bred in standard vivarium conditions and experimental procedures were approved by the Institutional Animal Care and Use Committee (IACUC) of the University of Utah. The *Stau1^tm1Apa(−/−)^* (*Stau1^−/−^*) mouse(5) was a generous gift from Prof. Michael A. Kiebler, Ludwig Maximilian University of Munich, Germany. *Stau1^−/−^* mice were maintained in a C57BL/6BJ background and Pcp2-ATXN2[Q127] (*ATXN2^Q127^*) mice^(21)^ were maintained in a B6D2F1/J background. *ATXN2^Q127^* (Pcp2-ATXN2[q127]) mice^22^ were crossed with *Stau1^tm1Apa(−/−)^* (*Stau1^−/−^*) mouse to generate *ATXN2^Q127/Tg^ Stau1^tm1Apa(+/−)^* and *ATXN2^Q127/Wt^ Stau1^tm1Apa(+/−^*). These mice were then bred to produce *ATXN2^Q127/Tg^ Stau1^tm1Apa(−/−)^* and *ATXN2^Q127/Wt^ Stau1^tm1Apa(−/−)^* in a mixed background of B6D2F1/J and C57BL/6J. Animals were genotyped according to previously published protocols (5, 21).

### Primary cultures of cortical neurons

Cultures of cortical neurons were prepared from WT or *Stau1^−/−^* neonatal mice euthanized according to IACUC approved protocols. Cortices from 6-7 animals were isolated, cut into 2mm segments and incubated with 50 units of papain (Worthington Biochemical, USA) in Earle’s balanced salt solution with 1. 0mM L-cysteine and 0.5mM EDTA for 15 minutes at 37°C. Digested tissue was washed with EBSS and mechanical dissociation was performed with a 1ml micropipette in the presence of 0.1mg/ml of DNase1 (Sigma-Aldrich). Cell suspension was filtered through a 40μm strainer (Corning) to remove any remaining aggregates. Neurons were seeded on poly-L-ornithine (Sigma-Aldrich) and laminin coated plates at a density of 50,000 per cm^2^ in Neurobasal Plus medium containing with 2% B27 Plus supplement. To prevent proliferation of glial cells 1μM cytosine arabinoside (Sigma-Aldrich) was added on day 2 and 90% of the media volume was changed after 24hrs. From there on, 60% of culture medium was replenished every 2-3 days. Experiments were conducted on day 9-10 by replacing all culture media with fresh media containing thapsigargin or vehicle (DMSO).

### DNA constructs, siRNA, cell treatments and transfections

Plasmid constructs 3xFlag-tagged STAU1 (3xF-STAU1) was prepared as detailed previously (2). All constructs were cloned into a pCMV-3xFlag plasmid (Agilent Technologies, USA) and verified by sequencing. The sequences or commercial origin of all siRNAs used in this study are listed in the supplementary table 4. For siRNA experiments HEK293 cells or fibroblasts were transfected with lipofectamine 2000 and harvested after 72hrs. For overexpression of recombinant proteins in HEK293 cells we utilized lipofectamine 3000 for 4hrs and harvested after 72hrs. For experiments involving both over-expression of recombinant protein and siRNA, HEK293 cells were transfected as specified for recombinant protein and 24hrs later for siRNA. Cells harvested 48hrs after the last transfection. Information on the pharmacological agents used to treat cells are listed in suppmentary table 3.

### Western blotting

Protein homogenates from cultured cells were prepared by scraping cells in phosphate buffered saline and lysing the pellets in Laemmli sample buffer (Bio-Rad), followed by boiling for 5 minutes. Tissues were manually homogenized with a pestle in extraction buffer (25 mM Tris-HCl pH 7.6, 300 mM NaCl, 0. 5% Nonidet P-40, 2 mM EDTA, 2 mM MgCl_2_, 0.5 M urea and protease inhibitors (Sigma-Aldrich)). After clarification supernatants were mixed with Laemmli buffer and boiled for 5 minutes. All protein extracts were resolved by SDS-PAGE and transferred to Hybond P membrane (Amersham Bioscience), blocked in Tris-buffered saline 0.1% Tween-20 with 5% skim milk and primary antibody was incubated overnight in this same solution or 5% bovine serum albumin when antibodies were directed against phosphorylated epitopes. Information about all antibodies used in supplementary table 2. After incubation with the corresponding secondary antibody signal was detected using Immobilon Western Chemiluminescent HRP Substrate (EMD Millipore) or SuperSignal™ West Pico PLUS Chemiluminescent Substrate (ThermoFisher Scientific) and photographed with a Bio-Rad ChemiDoc. Analysis and quantification was performed with Image Lab software (Bio-Rad). Relative protein abundance was first normalized against actin band intensity and then expressed as the ratio to the normalized control.

### Immunocytochemistry

Cells were cultured in NUNC Lab-Tek chamber slides, fixed with paraformaldehyde 4% in phosphate buffered saline and staining was performed according to previously published protocols (22, 23). Imaging was performed at the Fluorescence Microscopy Core Facility, a part of the Health Sciences Cores at the University of Utah.

### Quantitative RT-PCR

RNA extraction from cell cultures was performed with the RNeasy mini kit according to manufacturer’s instructions (Qiagen) and cDNA was prepared from 1ug of RNA with the ProtoScript cDNA synthesis kit (New England Biolabs). Quantitative RT-PCR was performed at the Genomics Core Facility, a part of the Health Sciences Cores at the University of Utah. PCR reactions were carried out with Sybr Green PCR Master mix (ThermoFisher Scientific). Primer sequences are listed in supplementary table 4. Gene expression was normalized to GAPDH levels and analyzed with the relative standard curve method.

### Statistical analysis

All results are presented as mean ± standard deviation unless noted otherwise. Comparisons between groups were made using the Student’s t Test in OriginPro 2017 software. Level of significance was set at P≤0.05. Levels of significance are noted as *P□≤0.05, **P□≤□10.01 and ns□=□P□>□0.05.

## Results

### ER stress causes increase in STAU1 mRNA and protein levels

To investigate STAU1 response to ER stress, we treated HEK293 cells with thapsigargin, which depletes ER calcium reserves, tunicamycin to inhibit N-linked glycosylation, ionomycin, a calcium ionophore, and brefeldin A, blocking secretion from the Golgi apparatus(24) and evaluated STAU1 levels and activation of the UPR after 18 hrs. Increasing doses of thapsigargin, tunicamycin, ionomycin and brefeldin A resulted in increasing abundances of both STAU1 and CHOP (Fig.1 a). The effects of thapsigargin on STAU1 mRNA and protein levels were apparent between 4 to 8 hrs after treatment (Fig.1 b, c).

**Fig. 1.**
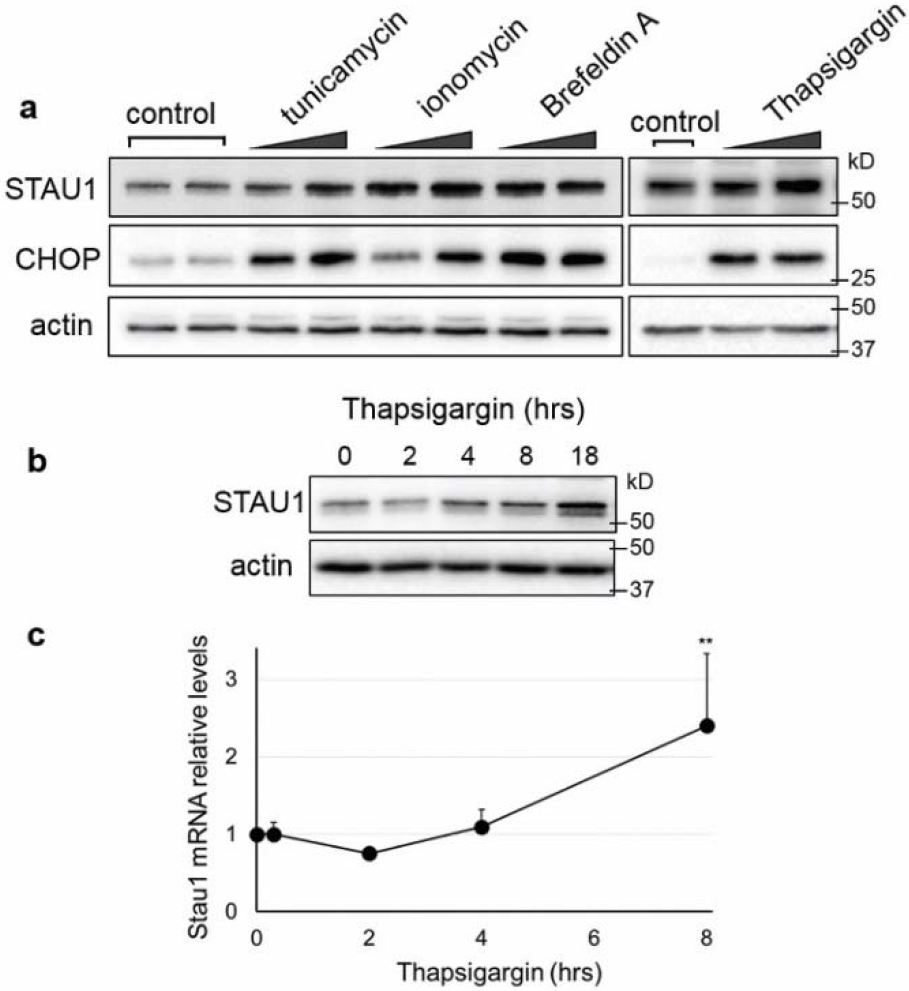
STAU1 overabundance induced by ER stress or calcium dyshomeostasis. **(a)** HEK293 cells were incubated with tunicamycin (0.1 and 0.5 μM), ionomycin (0.5 and 1 μM), Brefeldin A (0.5 and 1 μM) and thapsigargin (0.5 and 1 μM) for 18 hours. Levels of STAU1 and CHOP were evaluated by western blot. **(b)** Levels of STAU1 protein in HEK293 cells treated with thapsigargin (0.5 μM) for the times indicated **(c)** Relative STAU1 mRNA levels in HEK293 cells treated with thapsigargin (0.5 μM) for the times indicated. Data are mean□±□SD. **p < 0.01 according to paired sample t test.

### Lowering STAU1 protects from apoptosis induced by ER stress

In order to study the functional consequences of the STAU1 overabundance caused by ER stress in neurological disease, we examined the response of mouse primary cortical neurons and skin fibroblasts derived from WT, STAU1^+/−^ or STAU1^−/−^ animals to thapsigargin. In agreement with the results in Fig 1, thapsigargin elicited an increase in STAU1 in WT cells, along with large increases in CHOP and cleaved caspase 3. In STAU1^−/−^ neurons CHOP induction was significantly lower, and cleavage of caspase 3 was completely abolished (Fig. 2a). Similar results were seen in fibroblasts, where STAU^+/−^ cultures evidence a STAU1 dose-dependency of CHOP and cleaved caspase 3 activation (Fig. 2b).

**Fig. 2.**
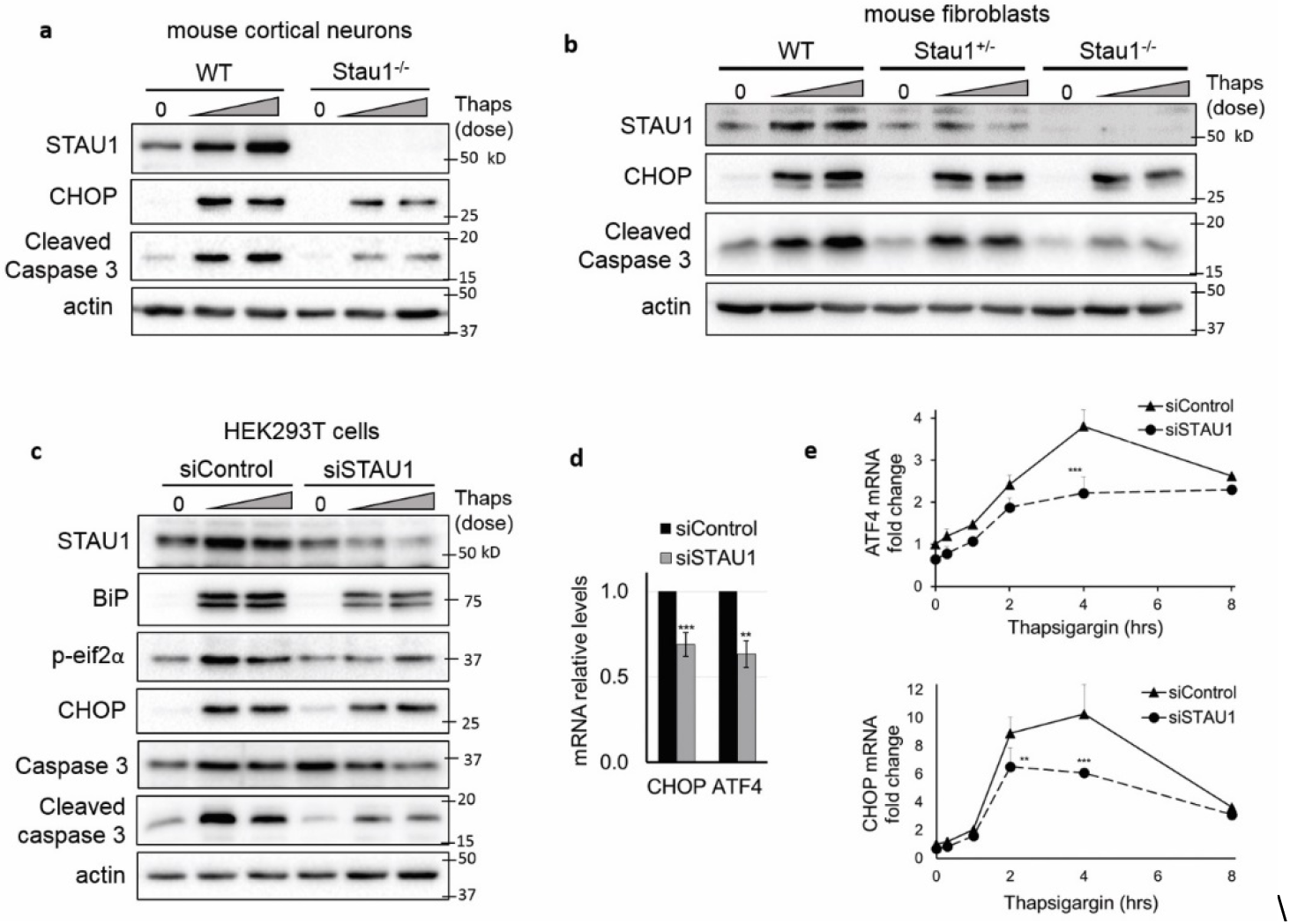
Attenuated UPR and apoptosis in cells deficient in STAU1. **(a and b)** western blots of cultured cortical neurons (a) or skin fibroblasts (b) from WT, STAU+/− or STAU1−/− mice incubated with thapsigargin (0.25 and 0.5 μM, 18 hrs). (c) western blots of HEK293 cells transfected with siControl or siSTAU1 for 72hrs and incubated with thapsigargin (0.5 and 1 μM) for 18 hrs. (d) mRNA levels of CHOP and ATF4 in HEK293 72 hours post-transfection with siControl or siSTAU1 and (e) after treatment with thapsigargin (0.5 μM) for the times indicated. Data are mean□±□SD. *p < 0.05, **p < 0.01 according to paired sample t test.

To further demonstrate the role of STAU1 in ER stress-induced apoptosis, HEK293 cells were transfected with a control siRNA (siControl) or a STAU1 siRNA (siSTAU1) and were challenged with thapsigargin for 18 hours. This compound caused a sharp induction of the UPR and apoptosis in control cells, whereas the response was greatly attenuated upon silencing of STAU1, characterized by lower levels of BiP, P-eIF2α, CHOP and cleaved caspase 3 (Fig. 2c).

To confirm that the role of STAU1 in ER stress-induced apoptosis was general to ER stress and not specific to thapsigargin, we analyzed cells treated with tunicamcycin or brefeldin A, which induce ER stress by vastly different mechanisms. We found STAU1 knockdown also attenuated the UPR and apoptotosis (Supplementary Fig. 1), indicating STAU1 has a role modulating life and death decisions when cells are faced with ER stress.

STAU1 silencing significantly reduced baseline levels of ATF4 and CHOP mRNA in HEK293 cells (Fig. 2d), and attenuated their induction by thapsigargin (Fig. 2e). Baseline levels of apoptotic factors were also significantly decreased after STAU1 silencing (Supplementary Fig.2b). Their transcriptional induction after thapsigargin did not reach statistical significance (not shown), in agreement with previous reports showing their acute activity is regulated mainly post-transcriptionally.

ATF4 and CHOP transcription increased immediately after addition of thapsigargin and peaked at 4 hours (Fig. 2e). In contrast, STAU1 protein and mRNA levels showed a delayed increase, only evident 4 to 8 hours after addition of thapsigargin (Fig. 1b, c and supplementary Fig. 2a). The fact that silencing STAU1 decreased both basal and induced levels of ATF4 and CHOP mRNAs even before overabundance of STAU1 was evident suggests that baseline levels of STAU1 may play a role in the modulation of ATF4 and CHOP mRNA levels, therefore overabundance of STAU1 might not be necessary to mediate its pro-apoptotic effects.

In all, our data show that STAU1 amplifies the activation of the UPR in a pro-apoptotic manner and knockdown or knockout of STAU1 is sufficient to prevent apoptosis during ER stress.

### STAU1 causes ER stress and apoptosis through the PERK-CHOP pathway

To understand the pathways by which STAU1 can modulate ER stress-induced apoptosis we studied cells expressing exogenous STAU1 in the absence of any other stressors or disease-related mutations. Exogenous STAU1 expression caused a substantial increase in the eIF2α kinase PERK, P-eIF2α and activation of caspase 3. This was prevented by a PERK inhibitor or siRNA against PERK (Fig 3a). These results indicate increased STAU1 signals through the PERK pathway of the UPR to cause apoptosis, and inhibiting this pathway was sufficient to completely prevent the effects of STAU1.

**Fig. 3.**
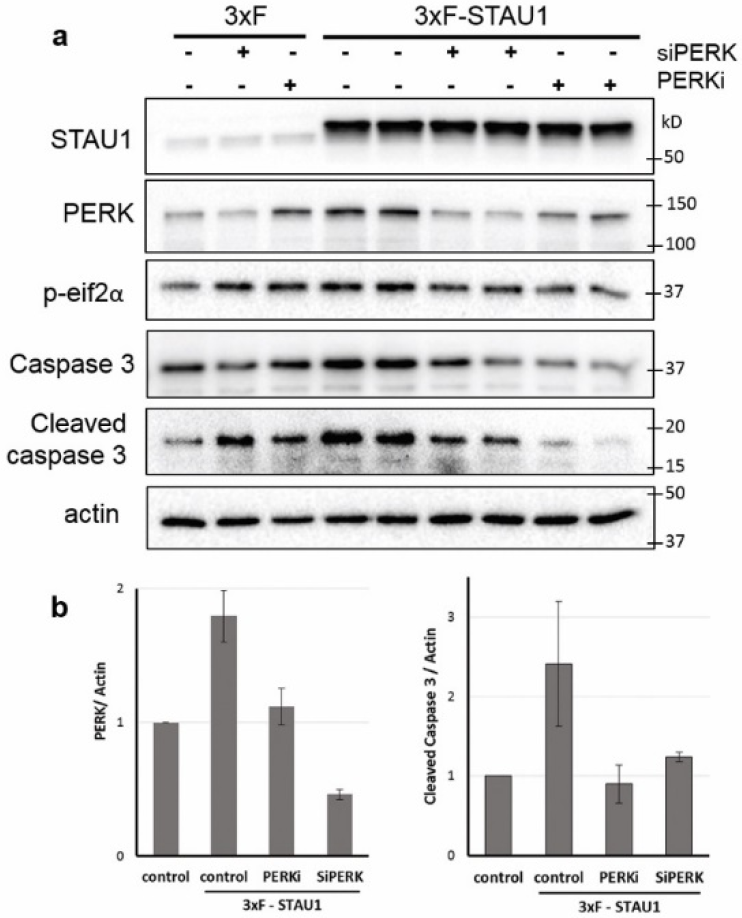
Exogenous STAU1 induces apoptosis through the PERK pathway of the UPR. **(a)** HEK293 cells were transfected with 3xFlag-STAU1 (3xF-STAU1) or empty vector control (3XF) with addition of siRNA directed at PERK (siPERK) after 24 hrs or the PERK inhibitor GSK2606414 (0.5 μM) after 48 hrs. After a further 18 hrs, protein levels were analyzed by western blot. **(b)** Quantification of three independent experiments. Data are mean□±□SD. *p < 0.05, **p < 0.01 according to paired sample t test.

Because STAU1 interacts with the eIF2α kinases PKR and CGN2 (25, 26), we studied all four eIF2α kinases for involvement in STAU1-mediated apoptosis. We found that each of them contributed to phosphorylation of eIF2α, but only PERK and PKR mediated apoptosis (supplementary Fig.3a). These results indicate that P-eIF2α is not required for apoptosis triggered by STAU1. In support of this, we saw increased apoptosis upon treatment with ISRIB, which prevents P-eIF2α effects without modifying its levels (27, 28) (Supplementary Fig. 3b). These results suggest that translational attenuation mediated by P-eIF2α is not required for apoptosis and also serves a protective role, indicating a complex interplay of various components of the integrated stress response (ISR).

### Decreasing STAU1 prevents ER stress and apoptosis in cellular and mouse models of SCA2

To study if the pro-apoptotic effects of STAU1 occured in cells stressed by mutations known to cause neurodegeneration, we studied cells and mice with mutations in ATXN2, the protein mutated in Spinocerebellar Ataxia type-2 (SCA2)(29). Previously we found calcium dyshomeostasis and increased STAU1 levels in SCA2 cells and mice (2, 30, 31) but we did not study whether this was associated with ER stress. We utilized HEK293 cells edited by CRISPR/Cas9 to introduce an expansion of 58 CAG repeats into one ATXN2 allele (ATXN2-Q58 cells) and the parental HEK293 cell line as control (ATXN2-Q22)(2). We found that ATXN2-Q58 cells had increased levels of the UPR proteins BiP, IRE1, P-eIF2α, spliced XBP1 and CHOP (Supplementary Fig. 4a), along with migration of CHOP to the nucleus (Supplementary Fig. 4b), hallmark of its active status. Activation of the UPR was a direct consequence of mutated ATXN2, as its silencing was sufficient to restore UPR mediators to levels comparable to normal cells (Supplementary Fig. 4a). Along with this, we found aggregates strongly immunoreactive for ubiquitin in ATXN2-Q58 cells, evidencing general disruption of cellular protein homeostasis (Supplementary Fig.4b).

Mouse models of SCA2 display increased STAU1 levels in the nervous system, and decreasing STAU1 protects Purkinje neurons and delays the onset of motor symptoms(2). The cellular mechanisms underlying this protection, however, are not fully understood. We studied whether STAU1 linked ATXN2 to apoptosis in models of SCA2. In ATXN2-Q58 cells, STAU1 silencing was sufficient to decrease UPR activation, evidenced by a significant reduction in the levels of BiP, IRE1, PERK, P-eIF2α and CHOP (Fig. 4a). In contrast, abundance of unspliced XBP1 and spliced XBP1 was increased by STAU1 silencing, likely in restorative effort, as XBP1 is essential to prevent cell death caused by ER stress (Fig. 4a). STAU1 silencing significantly decreased CHOP, total caspase 3 and cleaved caspase 3, indicating that STAU1 is necessary for pro-apoptotic activation of the UPR in this model (Fig.4a and b).

**Fig. 4.**
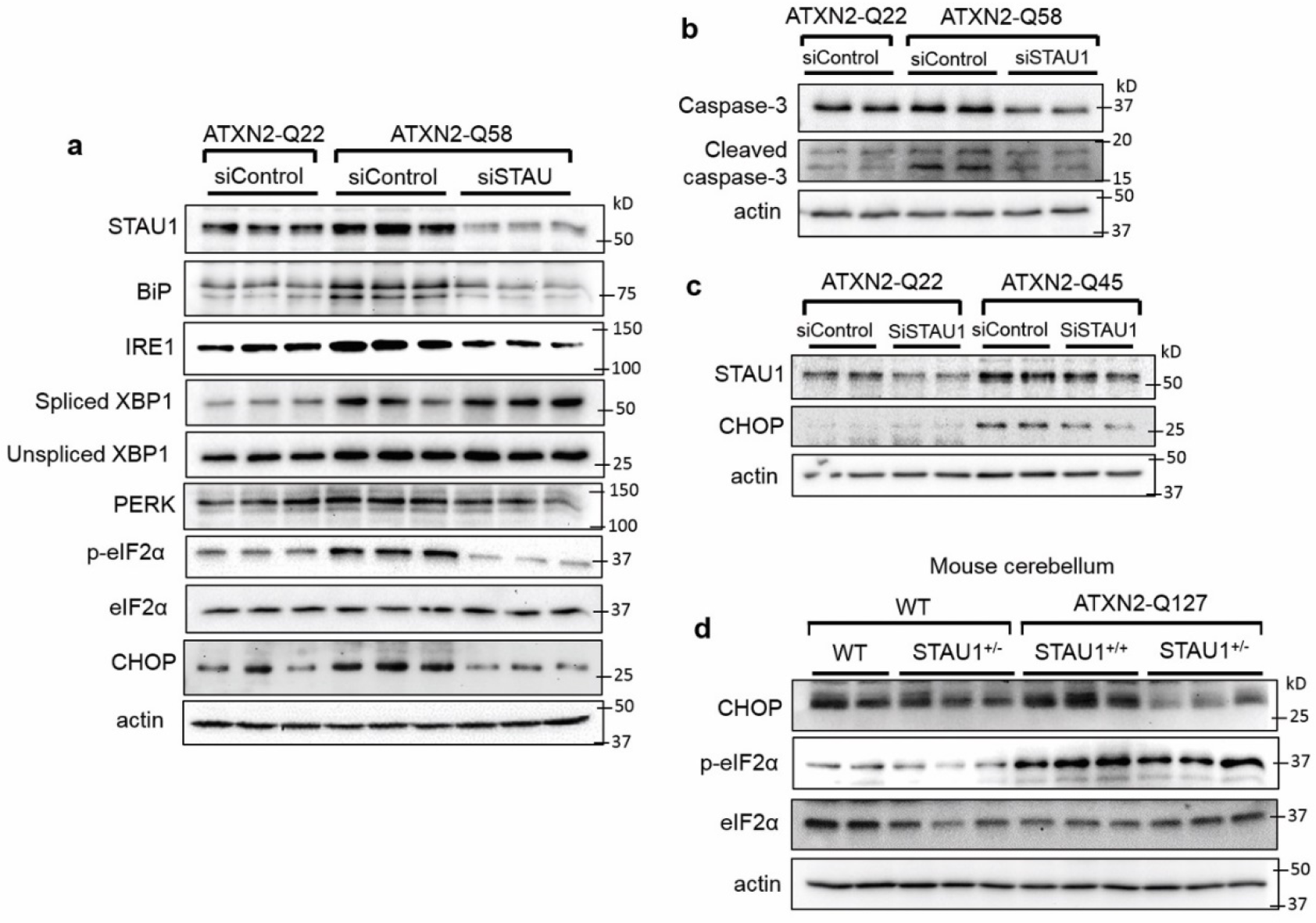
Attenuation of UPR and apoptosis in cellular and animal models of SCA2 by siSTAU1 or genetic interaction. **(a)** Western blot of proteins involved in the UPR, and **(b)** caspase 3 and cleaved caspase 3 in ATXN2-Q22 and ATXN2-Q58 cells **(c)** western blot of fibroblasts derived from an SCA2 patient with ATXN2-Q45 mutation. **(d)** Western blot of cerebellar tissue from WT, *ATXN2-Q127* mice and *Stau1^+/−^* haploinsufficient littermates at 34 weeks of age.

Fibroblasts derived from an SCA2 patient with a pathological ATXN2 expansion (ATXN2-Q45) recapitulated these findings, as evidenced by a significant decrease in CHOP when STAU1 was silenced (Fig. 4c). Increased CHOP and P-eIF2α in cerebella of ATXN2-Q127 mice was improved by STAU1 haploinsufficiency, indicating STAU1 can also modulate ER-stress induced apoptosis *in vivo* (Fig. 4d).

Because alterations in calcium homeostasis have been previously described in SCA2 models (30–34), we investigated whether they had a role in induction of apoptosis by STAU1 in ATXN2-Q58 cells. We found that STAU1 levels in ATXN2-Q58 cells were sensitive to changes in cytoplasmic calcium, as they were normalized by the intracellular calcium chelator BAPTA-AM and a CAMKK2 inhibitor (STO-609) (supplementary Fig. 5a and b). CAMKK2 is Ca^2+^/calmodulin-dependent protein kinase kinase that is activated in response to an increase in the cytosolic-free calcium. In addition, depleting IP3 levels with lithium and valproic acid or blocking calcium efflux from the ER with IP3R or RyR channel blockers (Xestospongin C, 2-ABP, dantrolene or DHBP) also decreased STAU1, indicating that both types of channels are involved in raising cytoplasmic calcium in ATXN2-Q58 cells (Supplementary Fig. 5a and b). These results provide evidence that a pro-apoptotic signaling axis involving calcium alterations, STAU1 and ER stress is active in this model of SCA2.

### Decreasing STAU1 prevents ER stress in cellular models of ALS and FTD

We have previously shown that STAU1 is increased in HEK293 cells transfected with WT TDP-43 or mutated TDP-43(2). Here we studied fibroblasts derived from two patients with mutations in the TDP-43 gene and two with expansions in C9ORF72, causative of ALS and FTD respectively. We found increased STAU1 along with a basally activated UPR in all of these patient cells, and silencing STAU1 was able to prevent UPR activation, including a strong decrease in CHOP, indicating STAU1 contributed to the pathological phenotype in these ALS and FTD models (Fig. 5 a and b)

**Fig. 5.**
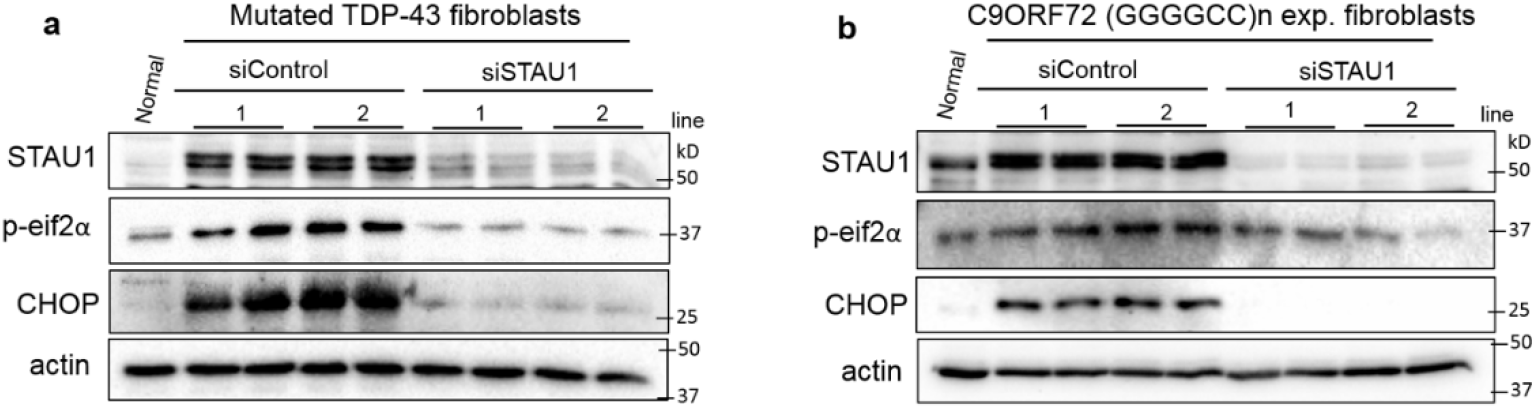
STAU1 knockdown reduces the pro-apoptotic activation of the UPR in fibroblasts from patients with ALS and FTD-causing mutations. **(a)** Western blot of STAU1, P-eIF2α and CHOP in fibroblasts derived from human subjects without disease-related mutation (normal), with TARDBP A382T (line 1) and one with TARDBP G298S (line 2) **(b)** and 2 individuals with C9ORF72 GGGGCC repeat expansion (line 1 and 2).

## Discussion

STAU1 is an RNA-binding protein with key roles in RNA metabolism and stress granule formation (2, 11). Previous reports highlight a striking overabundance of STAU1 in multiple models of neurological disease (2), however the functional consequences of this have not been fully explored. We have identified STAU1 as a modulator of apoptotic signaling during ER stress in multiple models of neurological disease and also in normal cells exposed to pharmacological stressors. This view is substantiated by a number of observations, namely, **(a)** ectopic expression of exogenous STAU1 caused apoptosis through the PERK-CHOP pathway **(b)** STAU1 knockout or knockdown cells showed attenuated UPR and apoptosis in response to ER stressors **(c)** Basal levels of UPR activation and apoptosis in cellular and mouse models of SCA2 and TDP-43 ALS and C9ORF72 FTD were markedly decreased by STAU1 knockdown.

Our data suggest that STAU1 lies both upstream and downstream of UPR activation. Ectopic overabundance of STAU1 was sufficient to induce ER stress, terminally activating the PERK-CHOP pathway (Fig. 3). Analogously, ER stress triggered STAU1 increase, creating a convergent maladaptive feed forward mechanism that amplified STAU1 abundance, ER stress and apoptosis (Fig. 1). The PERK-CHOP arm of the UPR is the canonical pro-apoptotic pathway, orchestrating cell death by inhibiting autophagy, increasing stress granule formation, altering the redox state of the cell, promoting expression of GADD45 (growth arrest and DNA-damage-inducible protein) and downregulating the anti-apoptotic mitochondrial protein BCL-2 (15, 35–38). These events lead to mitochondrial damage, release of cytochrome c and activation of caspase 3. In the present study we found that STAU1 could modulate the PERK pathway, upregulating ATF4 and CHOP and therefore precipitating cell death (Fig. 2 and 3).

Lack of caspase 3 cleavage was consistent with resistance to ER stress-induced apoptosis in STAU1 knockout and knockdown cells (Fig. 2a, b and c). With continued presence of the stressor, however, cell death was also noted in these cells. Possibly, at late phases of the stimulation excessive cellular damage activates additional non-apoptotic death mechanisms, and targeting the apoptotic pathway may not be enough to suppress cell death. Of note, in cells expressing mutant ATXN2, knockdown of STAU1 decreased activation of the three arms of the UPR, however it caused an increase in spliced XBP1 (Fig. 4a). This supports the notion that reducing STAU1 could protect the cells in two ways, by decreasing pro-apoptotic signaling and by increasing expression of adaptive and restorative factors such as spliced XBP1. XBP1 is recognized for conferring cancer cells chemoprotection, and also improving neurodegenerative diseases such as Alzheimer’s disease, Parkinson’s disease, spinal cord injury and ALS (14, 39–44).

We showed profound changes in eIF2α phosphorylation levels in response to STAU1 overexpression (Fig. 3) or silencing (Fig. 4 and 5) and in STAU1 knockout cells (Fig. 2). A previous report showed that modulating STAU1 abundance didn’t impact levels of phosphorylated eIF2α under normal or stress conditions, despite being able to alter stress granule dynamics (6). A probable reason for this discordance is that in Thomas et al P-eIF2α was analyzed a maximum of 3 hours after the addition of the stress, while we assessed it after 18 hrs of stimulation or in chronic pathological states generated by disease-causing mutations. Our results suggest that overabundance of STAU1, resulting in P-eIF2α elevations, could contribute to abnormal formation or persistence of stress granules, contributing to aberrant translation, ribostasis and proteostasis (1, 2, 18).

Our study of cells derived from patients with ATXN2, TDP-43 and C9ORF72 mutations, as well as in cerebella from SCA2 mice show that the STAU1-CHOP axis described here is basally active in all these neurodegeneration models (Fig. 4 and 5). We previously demonstrated that SCA2 mice benefit from STAU1 knockdown, with improvement of motor and molecular phenotypes and preservation of Purkinje cell firing frequency (2). Our results suggest that a decrease in ER stress and pro-apoptotic UPR could be responsible for these phenotypic improvements. Additionally, PERK is upregulated in several models of neurodegeneration, including overexpression of TDP-43, prion-related protein and tau, and its inhibition protects against neuronal damage (14, 45). These data support STAU1 as a preferred therapeutic target for neurological disease compared to PERK, since targeting PERK is limited by its pancreatic toxicity (46–49).

In conclusion, the present work sheds light into the role of STAU1 in multiple diseases with STAU1 overabundance by showing that it is a key modulator of ER-stress induced apoptosis. STAU1 overabundance caused by ER stress or calcium alterations reduces cellular resistance to ER stress and precipitates apoptosis through the PERK-CHOP pathway. By decreasing ER stress and specifically reducing P-eIF2α, targeting STAU1 could ameliorate proteostasis, ribostasis and aberrant stress granule phenotypes in diseases caused by ATXN2, TDP-43 and C9ORF72 mutations. Further understanding of the molecular mechanisms linking STAU1 and ER stress will provide insight needed to safely modulate death pathways for therapeutic benefit.

## Supporting information

STAU1 UPR paper supplemental figures and methods

## Acknowledgments

We thank Prof. Dr. Michael A. Kiebler, Ludwig Maximilian University of Munich, Germany for providing *Stau1^tm1Apa(−/−)^* (*Stau1^−/−^*) mice. We thank Prof. Monica Vetter, department of neurobiology and anatomy, and her laboratory staff for allowing the use of the Bio-Rad ChemiDoc, Dr. Clement Chow for insightful discussion of this project and Erika Aoyama and Brian Marshall for providing technical assistance. This work was supported by National Institutes of Neurological Disorders and Stroke (NINDS) grants R37NS033123 to SMP, R01NS097903 to DRS, and U01NS103883 and R21NS081182 to SMP and DRS. SMP was also supported by Target ALS, and the Noorda foundation.

## References

1. Paul S, Dansithong W, Gandelman M, Zu T, Ranum LPW, Figueroa KP, et al. Staufen blocks autophagy in neurodegeneration. bioRxiv. 2019:659649.

2. Paul S, Dansithong W, Figueroa KP, Scoles DR, Pulst SM. Staufen1 links RNA stress granules and autophagy in a model of neurodegeneration. Nat Commun. 2018;9(1):3648.

3. Liu J, Zhang KS, Hu B, Li SG, Li Q, Luo YP, et al. Systematic Analysis of RNA Regulatory Network in Rat Brain after Ischemic Stroke. Biomed Res Int. 2018;2018:8354350.

4. Krichevsky AM, Kosik KS. Neuronal RNA granules: a link between RNA localization and stimulation-dependent translation. Neuron. 2001;32(4):683–96.

5. Vessey JP, Macchi P, Stein JM, Mikl M, Hawker KN, Vogelsang P, et al. A loss of function allele for murine Staufen1 leads to impairment of dendritic Staufen1-RNP delivery and dendritic spine morphogenesis. Proc Natl Acad Sci USA. 2008;105(42):16374–9.

6. Thomas MG, Martinez Tosar LJ, Desbats MA, Leishman CC, Boccaccio GL. Mammalian Staufen 1 is recruited to stress granules and impairs their assembly. J Cell Sci. 2009;122(Pt 4):563–73.

7. Cho H, Kim KM, Han S, Choe J, Park SG, Choi SS, et al. Staufen1-mediated mRNA decay functions in adipogenesis. Mol Cell. 2012;46(4):495–506.

8. Gong C, Maquat LE. IncRNAs transactivate STAU1-mediated mRNA decay by duplexing with 3’ UTRs via Alu elements. Nature. 2011;470(7333):284–8.

9. Gong C, Kim YK, Woeller CF, Tang Y, Maquat LE. SMD and NMD are competitive pathways that contribute to myogenesis: effects on PAX3 and myogenin mRNAs. Genes Dev. 2009;23(1):54–66.

10. Dugre-Brisson S, Elvira G, Boulay K, Chatel-Chaix L, Mouland AJ, DesGroseillers L. Interaction of Staufen1 with the 5’ end of mRNA facilitates translation of these RNAs. Nucleic Acids Res. 2005;33(15):4797–812.

11. Kim YK, Furic L, Desgroseillers L, Maquat LE. Mammalian Staufen1 recruits Upf1 to specific mRNA 3’UTRs so as to elicit mRNA decay. Cell. 2005;120(2):195–208.

12. Ravel-Chapuis A, Belanger G, Yadava RS, Mahadevan MS, DesGroseillers L, Cote J, et al. The RNA-binding protein Staufen1 is increased in DM1 skeletal muscle and promotes alternative pre-mRNA splicing. The Journal of cell biology. 2012;196(6):699–712.

13. Bonnet-Magnaval F, Philippe C, Van Den Berghe L, Prats H, Touriol C, Lacazette E. Hypoxia and ER stress promote Staufen1 expression through an alternative translation mechanism. Biochem Biophys Res Commun. 2016;479(2):365–71.

14. Hetz C, Chevet E, Harding HP. Targeting the unfolded protein response in disease. Nat Rev Drug Discov. 2013;12(9):703–19.

15. Tabas I, Ron D. Integrating the mechanisms of apoptosis induced by endoplasmic reticulum stress. Nat Cell Biol. 2011;13(3):184–90.

16. Urra H, Dufey E, Lisbona F, Rojas-Rivera D, Hetz C. When ER stress reaches a dead end. Biochim Biophys Acta. 2013;1833(12):3507–17.

17. Walter P, Ron D. The unfolded protein response: from stress pathway to homeostatic regulation. Science. 2011;334(6059):1081–6.

18. Mahboubi H, Stochaj U. Cytoplasmic stress granules: Dynamic modulators of cell signaling and disease. Biochim Biophys Acta Mol Basis Dis. 2017;1863(4):884–95.

19. Panas MD, Ivanov P, Anderson P. Mechanistic insights into mammalian stress granule dynamics. The Journal of cell biology. 2016;215(3):313–23.

20. Ran FA, Hsu PD, Wright J, Agarwala V, Scott DA, Zhang F. Genome engineering using the CRISPR-Cas9 system. Nat Protoc. 2013;8(11):2281–308.

21. Hansen ST, Meera P, Otis TS, Pulst SM. Changes in Purkinje cell firing and gene expression precede behavioral pathology in a mouse model of SCA2. Hum Mol Genet. 2013;22(2):271–83.

22. Gandelman M, Peluffo H, Beckman JS, Cassina P, Barbeito L. Extracellular ATP and the P2X7 receptor in astrocyte-mediated motor neuron death: implications for amyotrophic lateral sclerosis. J Neuroinflammation. 2010;7:33.

23. Gandelman M, Levy M, Cassina P, Barbeito L, Beckman JS. P2X7 receptor-induced death of motor neurons by a peroxynitrite/FAS-dependent pathway. J Neurochem. 2013;126(3):382–8.

24. Oslowski CM, Urano F. Measuring ER stress and the unfolded protein response using mammalian tissue culture system. Methods Enzymol. 2011;490:71–92.

25. Sugimoto Y, Vigilante A, Darbo E, Zirra A, Militti C, D’Ambrogio A, et al. hiCLIP reveals the in vivo atlas of mRNA secondary structures recognized by Staufen 1. Nature. 2015;519(7544):491–4.

26. Dixit U, Pandey AK, Mishra P, Sengupta A, Pandey VN. Staufen1 promotes HCV replication by inhibiting protein kinase R and transporting viral RNA to the site of translation and replication in the cells. Nucleic Acids Res. 2016;44(11):5271–87.

27. Sidrauski C, McGeachy AM, Ingolia NT, Walter P. The small molecule ISRIB reverses the effects of elF2alpha phosphorylation on translation and stress granule assembly. eLife. 2015;4.

28. Sidrauski C, Acosta-Alvear D, Khoutorsky A, Vedantham P, Hearn BR, Li H, et al. Pharmacological brake-release of mRNA translation enhances cognitive memory. eLife. 2013;2:e00498.

29. Pulst SM, Nechiporuk A, Nechiporuk T, Gispert S, Chen XN, Lopes-Cendes I, et al. Moderate expansion of a normally biallelic trinucleotide repeat in spinocerebellar ataxia type 2. Nat Genet. 1996;14(3):269–76.

30. Liu J, Tang TS, Tu H, Nelson O, Herndon E, Huynh DP, et al. Deranged calcium signaling and neurodegeneration in spinocerebellar ataxia type 2. J Neurosci. 2009;29(29):9148–62.

31. Kasumu AW, Liang X, Egorova P, Vorontsova D, Bezprozvanny I. Chronic suppression of inositol 1,4,5-triphosphate receptor-mediated calcium signaling in cerebellar purkinje cells alleviates pathological phenotype in spinocerebellar ataxia 2 mice. J Neurosci. 2012;32(37):12786–96.

32. Halbach MV, Gispert S, Stehning T, Damrath E, Walter M, Auburger G. Atxn2 Knockout and CAG42-Knock-in Cerebellum Shows Similarly Dysregulated Expression in Calcium Homeostasis Pathway. Cerebellum. 2016.

33. Kasumu A, Bezprozvanny I. Deranged calcium signaling in Purkinje cells and pathogenesis in spinocerebellar ataxia 2 (SCA2) and other ataxias. Cerebellum. 2012;11(3):630–9.

34. Bezprozvanny I. Role of inositol 1,4,5-trisphosphate receptors in pathogenesis of Huntington’s disease and spinocerebellar ataxias. Neurochem Res. 2011;36(7):1186–97.

35. Matsumoto M, Minami M, Takeda K, Sakao Y, Akira S. Ectopic expression of CHOP (GADD153) induces apoptosis in M1 myeloblastic leukemia cells. FEBS Lett. 1996;395(2–3):143–7.

36. Ron D, Habener JF. CHOP, a novel developmentally regulated nuclear protein that dimerizes with transcription factors C/EBP and LAP and functions as a dominant-negative inhibitor of gene transcription. Genes Dev. 1992;6(3):439–53.

37. Zinszner H, Kuroda M, Wang X, Batchvarova N, Lightfoot RT, Remotti H, et al. CHOP is implicated in programmed cell death in response to impaired function of the endoplasmic reticulum. Genes Dev. 1998;12(7):982–95.

38. McCullough KD, Martindale JL, Klotz LO, Aw TY, Holbrook NJ. Gadd153 sensitizes cells to endoplasmic reticulum stress by down-regulating Bcl2 and perturbing the cellular redox state. Mol Cell Biol. 2001;21(4):1249–59.

39. Gerakis Y, Hetz C. A decay of the adaptive capacity of the unfolded protein response exacerbates Alzheimer’s disease. Neurobiol Aging. 2018;63:162–4.

40. Cisse M, Duplan E, Lorivel T, Dunys J, Bauer C, Meckler X, et al. The transcription factor XBP1s restores hippocampal synaptic plasticity and memory by control of the Kalirin-7 pathway in Alzheimer model. Mol Psychiatry. 2017;22(11):1562–75.

41. Casas-Tinto S, Zhang Y, Sanchez-Garcia J, Gomez-Velazquez M, Rincon-Limas DE, Fernandez-Funez P. The ER stress factor XBP1s prevents amyloid-beta neurotoxicity. Hum Mol Genet. 2011;20(11):2144–60.

42. Loewen CA, Feany MB. The unfolded protein response protects from tau neurotoxicity in vivo. PLoS One. 2010;5(9).

43. Martinez A, Lopez N, Gonzalez C, Hetz C. Targeting of the unfolded protein response (UPR) as therapy for Parkinson’s disease. Biol Cell. 2019.

44. Valenzuela V, Collyer E, Armentano D, Parsons GB, Court FA, Hetz C. Activation of the unfolded protein response enhances motor recovery after spinal cord injury. Cell Death Dis. 2012;3:e272.

45. Rivas A, Vidal RL, Hetz C. Targeting the unfolded protein response for disease intervention. Expert Opin Ther Targets. 2015;19(9):1203–18.

46. Moreno JA, Halliday M, Molloy C, Radford H, Verity N, Axten JM, et al. Oral treatment targeting the unfolded protein response prevents neurodegeneration and clinical disease in prion-infected mice. Sci TransI Med. 2013;5(206):206ra138.

47. Halliday M, Radford H, Sekine Y, Moreno J, Verity N, le Quesne J, et al. Partial restoration of protein synthesis rates by the small molecule ISRIB prevents neurodegeneration without pancreatic toxicity. Cell Death Dis. 2015;6:e1672.

48. Yu Q, Zhao B, Gui J, Katlinski KV, Brice A, Gao Y, et al. Type I interferons mediate pancreatic toxicities of PERK inhibition. Proc Natl Acad Sci USA. 2015;112(50):15420–5.

49. Kaufman RJ, Back SH, Song B, Han J, Hassler J. The unfolded protein response is required to maintain the integrity of the endoplasmic reticulum, prevent oxidative stress and preserve differentiation in beta-cells. Diabetes Obes Metab. 2010;12 Suppl 2:99–107.

