## Supplementary material for "Staufen 1 amplifies pro-apoptotic activation of the unfolded protein response": STAU1 UPR paper supplemental figures and methods

**Supplementary figures and tables**

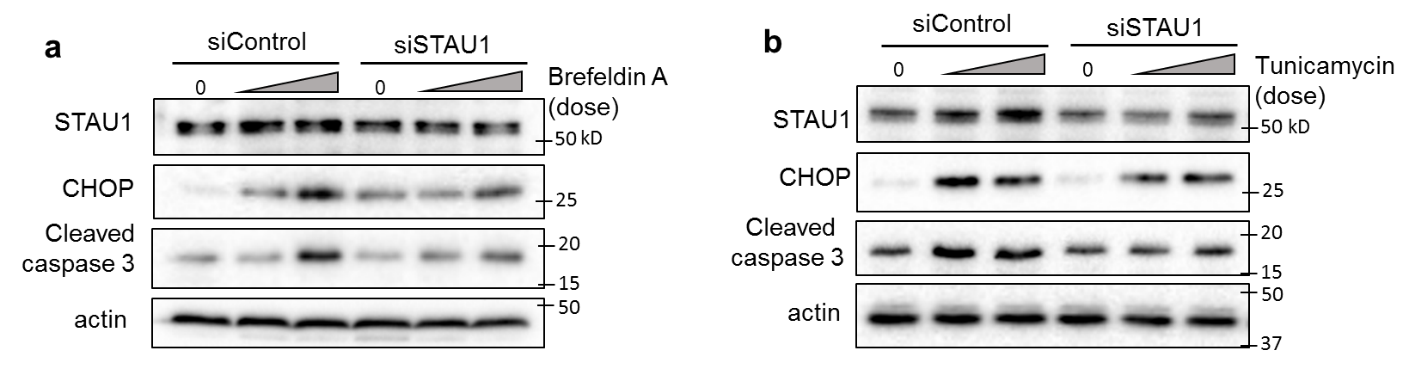

**Supplementary figure 1.** **STAU1 knockdown attenuates UPR and apoptosis caused by brefeldin A and tunicamycin.** Western blots of STAU1, CHOP and cleaved caspase 3 in HEK293 cells transfected with siControl or siSTAU1 and treated with **(A)** brefeldin A (1 µM) or **(B)** tunicamycin (1 µg/ml) after 72 hours.

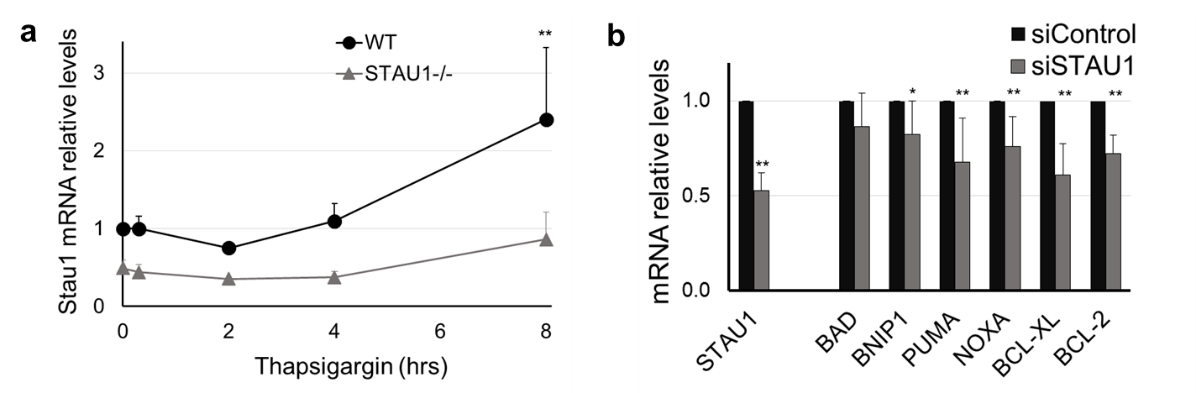

**Supplementary Fig. 2.** **(a)** Relative mRNA levels of Stau1 in HEK293 cells transfected with siControl or siSTAU1 and treated with thapsigargin after 72hrs. **(b)** Relative mRNA levels of apoptotic factors in HEK293 cells 72 hours after transfection with siControl or siSTAU1. Data are mean ± SD of at least 3 independent experiments. ^∗^p < 0.05, ^∗∗^p < 0.01; paired sample t test.

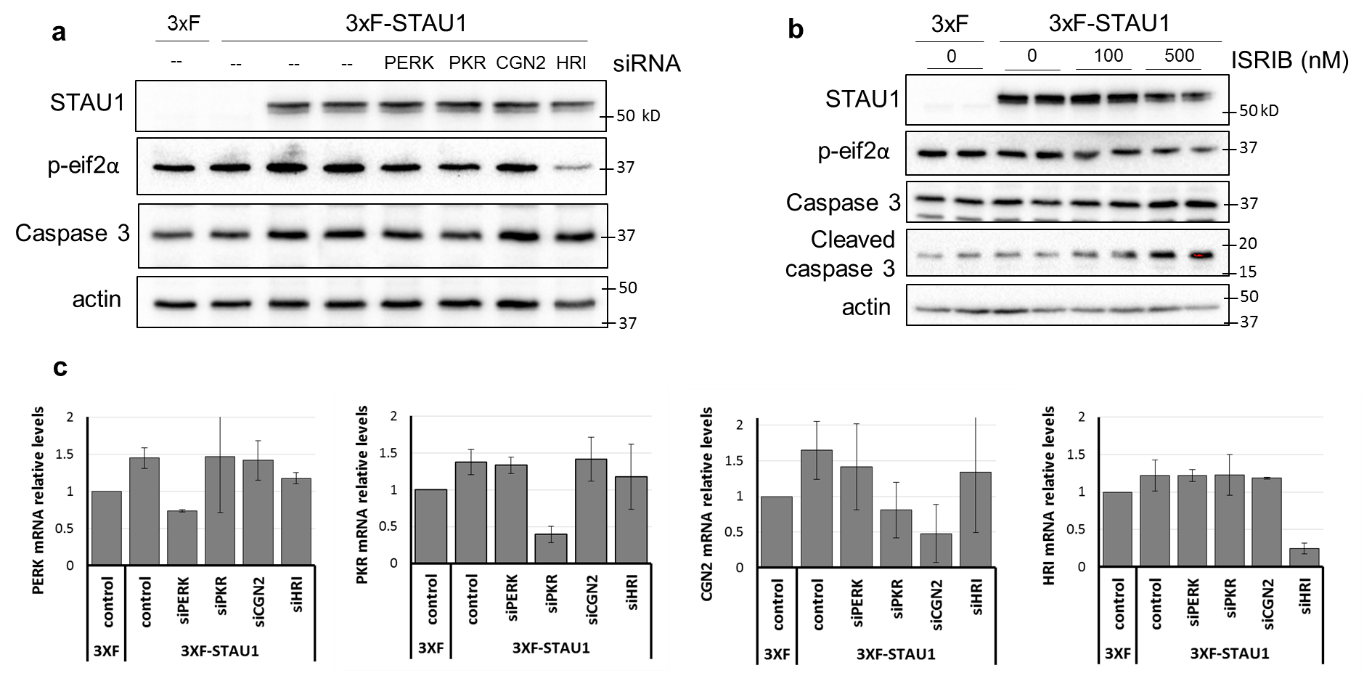
\

**Supplementary Fig. 3. Role of eif2 kinases and p-eIF2α in STAU1-mediated apoptosis**. **(a)** western blots of HEK293 cells transfected with 3xF or 3xF-STAU1, and subsequently with a siControl (--) or the indicated siRNA the day after. **(b)** Western blots of HEK293 cells transfected with 3xF or 3xF-STAU1 and 72 hours later treated with ISRIB (18 hrs) at the indicated concentrations. **(c)** Relative levels of the indicated genes after transfection as specified in (a).

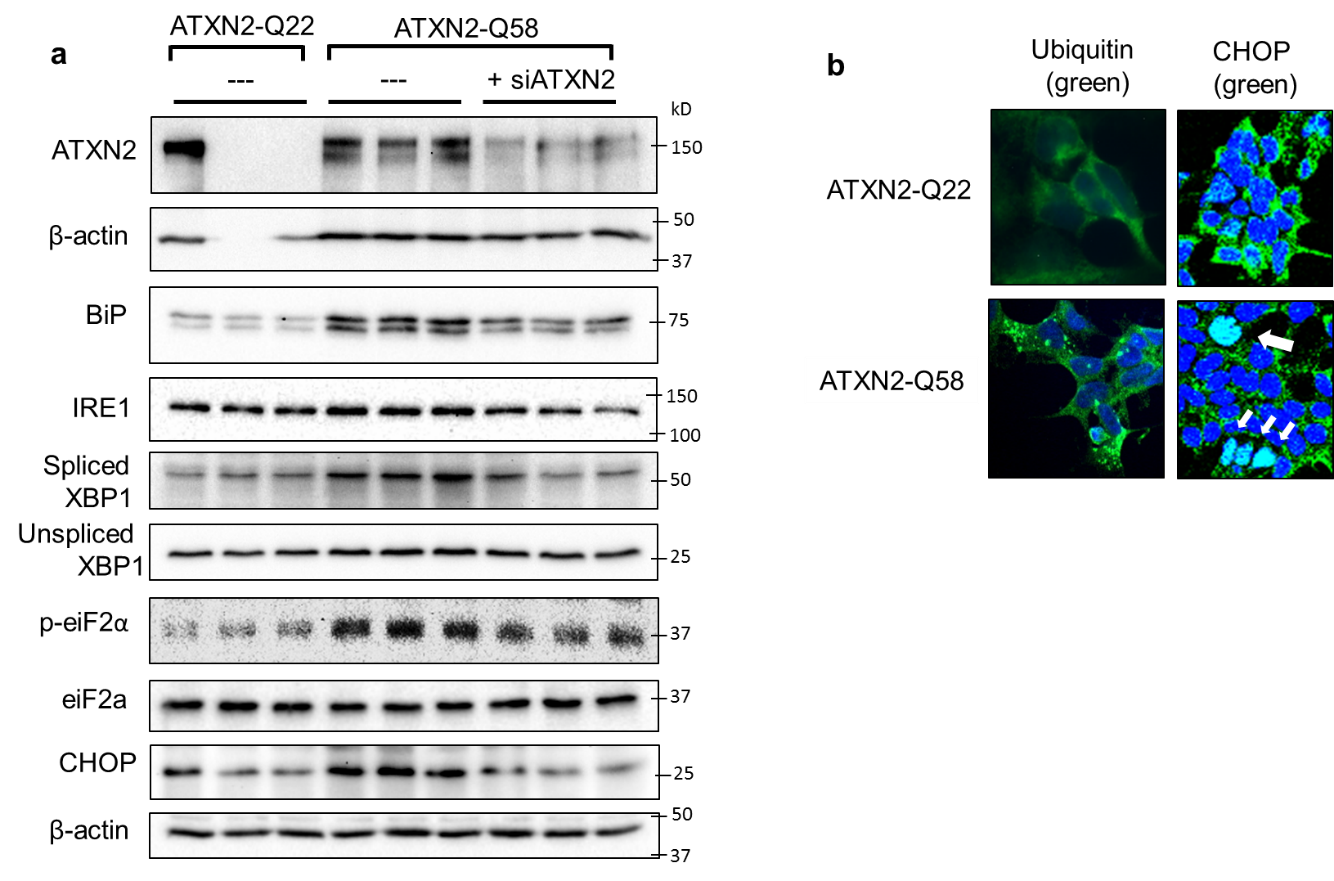

**Supplementary Fig. 4 ATXN2-dependent UPR activation, ubiquitinated aggregates and nuclear CHOP in ATXN2Q58 cells. (a)** Western blot of ATXN2 and UPR proteins in HEK293 cells expressing endogenous ATXN2Q22 or ATXN2Q58 cells and ATXN2Q58 cells with SiRNA for ATXN2 **(b)** Immunofluorescence for ubiquitin (green, left panels) and CHOP (green, right panels). Arrows indicate CHOP located in the cell nucleus.

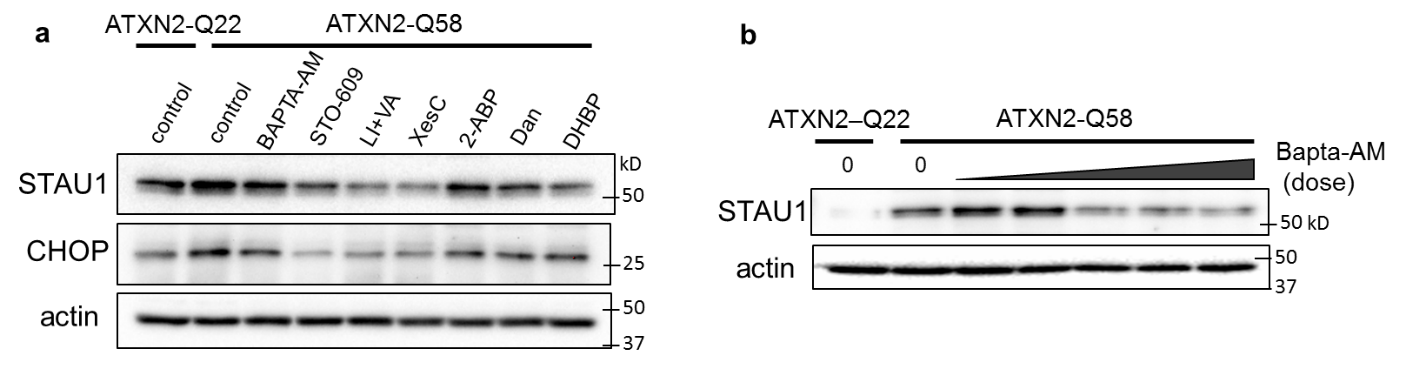

**Supplementary Fig. 5. STAU1 levels are modulated by calcium in ATXN2-Q58 cells (a)** Analysis of STAU1 levels in ATXN2-Q22 or ATXN2-Q58 cells after a 24 hour incubation with the calcium chelator BAPTA-AM (10 µM) CAMKK kinase inhibitor STO-609 (1 µM), IP3 depleting agents lithium + valproic acid (Li+VA, 1mM each), IP3 receptor inhibitors Xestospongin C (Xes C, 0.1 µM) and 2-ABP (1 µM), RyR inhibitors Dantrolene (Dan, 1 µM) and DHBP (1 µM). **(b)** STAU1 levels decrease in response to BAPTA-AM (1, 5, 10, 20 and 30 µM).

**Supplementary table 1**

| **Cell** | **SOURCE** | **IDENTIFIER** |
| --- | --- | --- |
| HEK293 | ATCC | CRL-1573 |
| HEK293-ATXN2-Q58 | Paul et al, 2018 | N/A |
| SCA2 fibroblasts – ATXN2-Q45 | Paul et al, 2018 | N/A |
| TDP-43 fibroblast line 1 - TARDBP p.Gly298Ser (c.892G>A) | NINDS human cell and data repository | ND32947 |
| TDP-43 fibroblast line 2 - TARDBP p.Ala382Thr (c.1144G>A) | NINDS human cell and data repository | ND41003 |
| C9orf72 fibroblast line 1 - GGGGCC repeat expansion; in 1st intron | NINDS human cell and data repository | ND42504 |
| C9orf72 fibroblast line 2 - GGGGCC repeat expansion; in 1st intron | NINDS human cell and data repository | ND42506 |
| Normal control patient | NINDS human cell and data repository | ND29510 |

**Supplementary table 2**

| **Antibody** | **SOURCE** | **IDENTIFIER** |
| --- | --- | --- |
| Mouse Anti-Ataxin-2 Clone  22 | BD Biosciences | 611378 |
| Staufen | Novus Biologicals | NBP1-33202 |
| CHOP (L63F7) Mouse mAb | Cell Signaling Technology | 2895 |
| eIF2α (L57A5) Mouse mAb | Cell Signaling Technology | 2103 |
| Phospho-eIF2α (Ser51) Antibody | Cell Signaling Technology | 9721 |
| PERK (C33E10) Rabbit mAb | Cell Signaling Technology | 3192 |
| IRE1α (14C10) Rabbit mAb #3294 | Cell Signaling Technology | 3294 |
| Cleaved Caspase-3 (Asp175) (5A1E) Rabbit mAb | Cell Signaling Technology | 9664 |
| Caspase 3 | ThermoFisher Scientific | PA5-16335 |
| Grp78 antibody (BIP) | GeneTex | GTX113340 |
| β-Actin−Peroxidase | Sigma-millipore | A3854 |
| Anti-XBP1 antibody | Abcam | ab37152 |
| Peroxidase-conjugated horse anti-mouse IgG | Vector Laboratories | PI-2000 |
| Peroxidase AffiniPure Goat Anti-Rabbit IgG | Jackson ImmunoResearch Laboratories | 111-035-144 |

**Supplementary table 3**

| **Chemical** | **SOURCE** | **IDENTIFIER** |
| --- | --- | --- |
| Thapsigargin | TOCRIS | 1138 |
| Ionomycin | TOCRIS | 1704 |
| DHBP | TOCRIS | 0839 |
| STO-609 | TOCRIS | 1551 |
| BAPTA-AM | TOCRIS | 2787 |
| ISRIB | TOCRIS | 5284 |
| Xestospongin C | TOCRIS | 1280 |
| Dantrolene | TOCRIS | 0507 |
| Tunicamycin | Cayman chemical | 11445 |
| Brefeldin A | Sigma-millipore | B6542 |
| Lithium chloride | Sigma-millipore | L4408 |
| Valproic acid | Stemgent | 04-0007 |
| 2-ABP | Calbiochem | 100065 |
| GSK260614 | APExBio | A3448 |

**Supplementary table 4**

| **Oligonucleotide** | **SOURCE** | | **IDENTIFIER** |
| --- | --- | --- | --- |
| Human primer Stau1 – forward - TCCTTGGTTTCAAAGTCCCG | | DNA/Peptide Facility, Health Sciences Center Cores, University of Utah. |  |
| Human primer BBC3 (PUMA) – forward -  ACCTCAACGCACAGTACGAG | | DNA/Peptide Facility |  |
| Human primer PMAIP1 (NOXA) – forward - TCCTGAGCAGAAGAGTTTGG | | DNA/Peptide Facility |  |
| Human primer BCL2 – forward - TTGCTTTACGTGGCCTGTTTC | | DNA/Peptide Facility |  |
| Human primer BCL-XL – forward - GTTCCCTTTCCTTCCATCC | | DNA/Peptide Facility |  |
| Human primer adenovirus E1B 19kDa interacting protein 1 (BNIP1) – forward - GTGTGAGGAGCTGCTGGCTTG | | DNA/Peptide Facility |  |
| Human primer BCL2-antagonist of cell death (BAD) – forward - AGGATCCGTGCTGTCTCCTTTG | | DNA/Peptide Facility |  |
| Human primer – DDIT3 (CHOP) – forward - AGAACCAGGAAACGGAAACAGA | | DNA/Peptide Facility |  |
| Human primer – ATF4 – forward - GTTCTCCAGCGACAAGGCTA | | DNA/Peptide Facility |  |
| Human primer eif2ak1 (HRI) – forward - CCACTTCGTTCAAGACAGGTG | | DNA/Peptide Facility |  |
| Human primer eif2ak2 (PKR) – forward - GGAAAGCGAACAAGGAGTAAGG | | DNA/Peptide Facility |  |
| Human primer eif2ak3 (PERK) – forward - GTCCGGAACCAGACGATGAG | | DNA/Peptide Facility |  |
| Human primer eif2ak4 (CGN2) – forward - GAAGCTGTCAGCCAGCACTA | | DNA/Peptide Facility |  |
| Human primer – GAPDH – forward -  CACATGGCCTCCAAGGAGTAA | | DNA/Peptide Facility |  |
| All Star Negative Control siRNA | | Qiagen | 1027280 |
| Human siATXN2 (Hs_ATXN2_2) | | Qiagen | SI00308196 |
| Human siSTAU1 siRNA – 5′CCUAUAACUACAACAUGAGdTdT-3′ | | Kim et al, 2005/ manufactured by ThermoFisher |  |
| Human siCHOP (Hs_DDIT3_1 FlexiTube siRNA) | | Qiagen | SI00059528 |
| Human siPKR (Hs_EIF2AK2_2 FlexiTube siRNA) | | Qiagen | SI00042819 |
| Human siCGN2 (Hs_EIF2AK4_1 FlexiTube siRNA) | | Qiagen | SI03054807 |
| Human siHRI (Hs_EIF2AK1_1 FlexiTube siRNA) | | Qiagen | SI00105784 |
| SignalSilence® PERK siRNA I | | Cell Signaling Technologies | 9024 |
